# Working memory and reasoning benefit from different modes of large-scale brain dynamics in healthy older adults

**DOI:** 10.1101/202630

**Authors:** Alexander V. Lebedev, Jonna Nilsson, Martin Lövdén

## Abstract

Researchers have proposed that solving complex reasoning problems, a key indicator of fluid intelligence, involves the same cognitive processes as solving working memory tasks. This proposal is supported by an overlap of the functional brain activations associated with the two types of tasks and by high correlations between inter-individual differences in performance. We replicated these findings in fifty-three older subjects but also showed that solving reasoning and working memory problems benefits from different configurations of the functional connectome and that this dissimilarity increases with higher difficulty load. Specifically, superior performance in a typical working memory paradigm (n-back) was associated with up-regulation of modularity (increased between-network segregation), whereas performance in the reasoning task was associated with effective down-regulation of modularity. We also showed that working memory training promotes task-invariant increases in modularity. Since superior reasoning performance is associated with down-regulation of modular dynamics, training may thus have fostered an inefficient way of solving the reasoning tasks. This could help explain why working memory training does little to promote complex reasoning performance. The study concludes that complex reasoning abilities cannot be reduced to working memory and suggests the need to reconsider the feasibility of using working memory training interventions to attempt to achieve effects that transfer to broader cognition.

## INTRODUCTION

Fluid intelligence is central to solving complex reasoning tasks; that is, to identifying common patterns and applying logic to extrapolate them to novel problems. The cognitive processes that determine performance in such tasks have been a matter of debate for decades (1-5). Task analyses have led some researchers to propose that reasoning ability depends on and is almost isomorphic with the capacity to maintain and manipulate information in working memory (5, 6). In keeping with this line of reasoning, psychologists have emphasized the demands that both working memory and reasoning tasks place on executive control (7, 8). The high correlation between inter-individual differences in working memory and complex reasoning scores (4, 5) and observations that these tasks elicit similar functional brain activity in the frontoparietal control network (CN) (8-14) are also consistent with this idea. Yet there are reasons to believe that cognitive processes other than working memory ability are involved in complex reasoning. Among them are exploratory cognitive functions such as perspective shifting and aspects of creativity that may facilitate rule discovery and insight. The latter refers to the spontaneous realization that a difficult problem has a simple solution if perceived from a conceptually different or new perspective. While these processes are much less studied, their importance in solving complex reasoning tasks has been demonstrated in psychometric studies (15, 16).

Functional brain imaging studies of large-scale brain dynamics allow us to disentangle some of the key principles underlying higher cognition. Shaped by evolution to enhance adaptability and efficient information processing, the brain appears to function as a critical system achieving optimal performance near phase transitions from order to disorder (17). This, in turn, requires a hierarchical modular structure balancing segregation and integration of its functional subdivisions (18-20). Modularity has been used as a global measure of such neurodynamics in studies of age-related changes (21, 22), working memory (23, 24), learning (25, 26), reactivity to environmental irregularities (27) and visual awareness (28). While task performance is generally associated with lower modularity than resting state (29-31), superior performance on cognitive tasks requiring *outward* attentional focus (e.g., working memory tasks) has been linked to relatively more modular (i.e., segregated) functional connectome at rest (23), whereas tasks requiring *inward* attentional focus, such as odor recognition and episodic memory retrieval, are instead linked to low-modular (i.e., more globally integrated) states (32, 33). Relatedly, effective suppression of the default mode network (DMN; functionally integrated group of mostly midline brain structures involved in self-referential processing and other tasks requiring inward focus) is related to performance on problems requiring outward attentional focus (34, 35). An opposite association has been reported for more complex tasks that demand creativity, cognitive flexibility, and, importantly, complex reasoning, in which resting state and task-related connectivity between the DMN and CN have been linked to superior performance (13, 36, 37). Furthermore, the direction of the association between modularity and performance may depend on task complexity: performance on simple tasks is positively associated with modularity, whereas performance on more complex tasks, in contrast, appears to be positively associated with low-modular states (38).

These disparate findings lead to the hypothesis that, despite a high correlation between performance and convergence of brain activation patterns, reasoning tasks require different, or at least additional, cognitive processes and different modes of large-scale brain dynamics than working memory. Better complex reasoning performance may be linked to a low-modular configuration of the functional connectome characterized by enhanced communication of the CN with the DMN and other brain networks. This enhanced communication may in turn promote an exploratory mode of cognition (39) – a state of “expected uncertainty” (40) – beneficial to rule discovery and abstraction (41). In contrast, successful solution of working memory tasks may require a more modular network configuration with stabilized configuration of the frontoparietal network coupled with effective suppression of the DMN activity to prevent inward attentional focus and mind-wandering. These divergent associations may be particularly marked at higher difficulty loads, because complex reasoning requires a more exploratory mode of cognition than simpler reasoning, one that benefits pattern discovery. It follows from these hypotheses that improvements from training working memory should lead to more modular brain dynamics during task performance. If such a shift in modularity generalizes to solving the reasoning task then this may only be beneficial for simple reasoning performance but not for solving the complex reasoning tasks that may require a higher degree of between-network integration. We employed large-scale network analysis to investigate these hypotheses. The study used functional magnetic resonance imaging (fMRI) during one reasoning and working memory task. The spatial reasoning task was developed to be similar to the classic matrix reasoning tests and a classical figural updating task (n-back) was used to probe working memory. Both tasks were performed at three different difficulties **(Figure 1**) in a sample of healthy older adults (*n* = 53; age = 65-75 years) before (i.e., at pretest) and after (i.e., at posttest) working memory training (n = 27) and active control training (perceptual matching; n = 26).

**Fig. 1.**
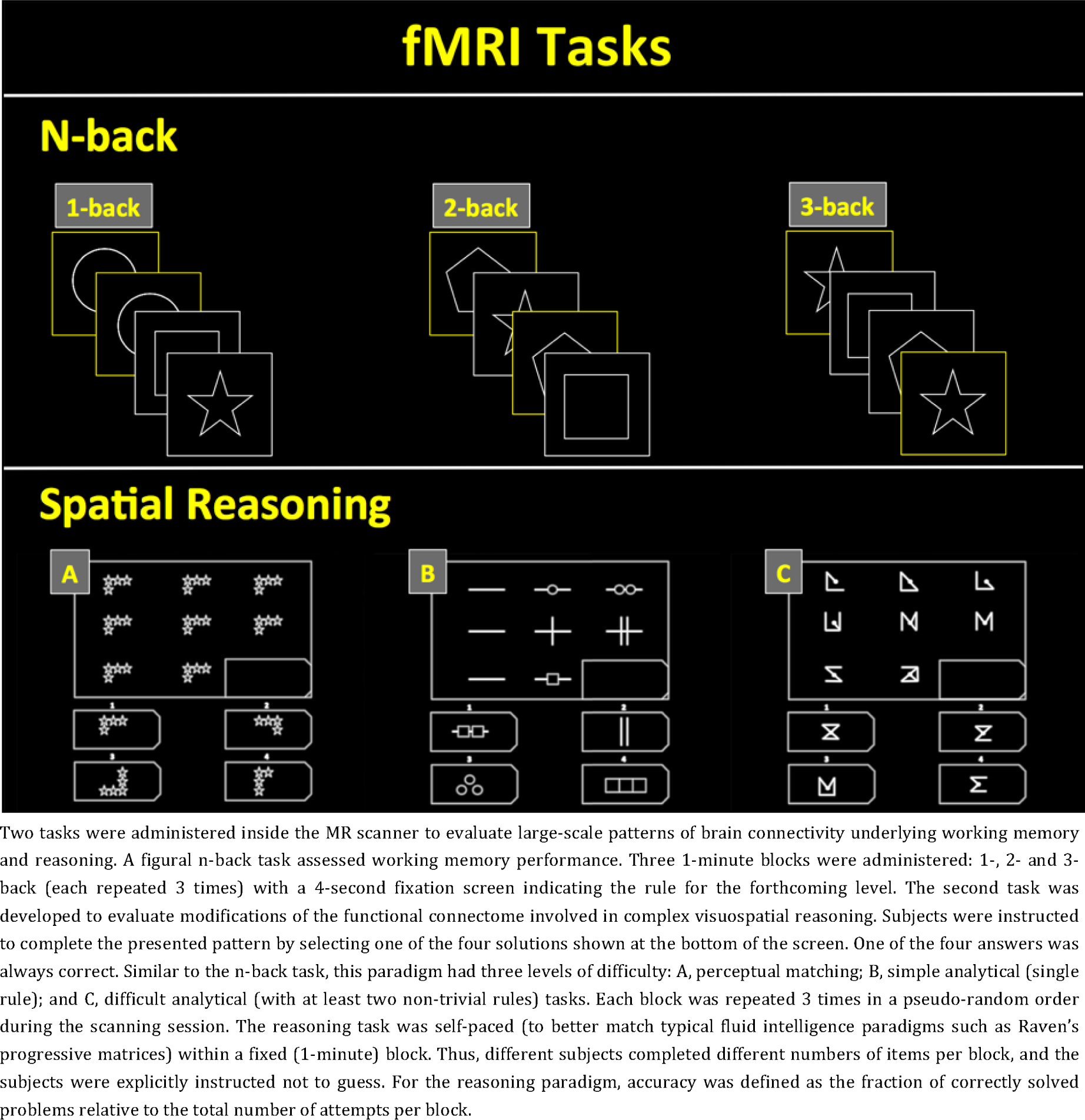
fMRI tasks.

## Methods

### Subjects

For the present analysis, the sample consisted of 53 participants who had brain scans of acceptable quality from two fMRI scanning sessions, one before and one after four weeks of cognitive training. Imaging data quality criteria were: i) no motion artifacts in the raw data noticeable to the naked eye, ii) max frame-wise displacement below 3mm, iii) less than 10% of missing volumes after the de-spiking procedure. The study was approved by the Regional Ethical Review Board in Stockholm (DNR: 2014/2188-31/1) and was conducted in accordance with the revised declaration of Helsinki [2013]. Informed consent was obtained from all subjects prior to enrollment. Standardized physical and neuropsychological examinations were also carried out at screening. The subjects were randomly assigned to one of the two alternative training programs: (1) working memory (active; *n* = 28; age = 68.89±3.13 years; education = 15.17±3.25 years; number of women = 14; attendance = 19.07±0.94 session) and (2) perceptual matching (control; *n* = 25; age = 69.48±2.96 years; education = 15.02±4 years; number of women = 14; Attendance = 19±1.58 session) and were blind to the hypotheses about the two protocols. The randomization was accomplished in an automated manner using R-programming language (42) for each study wave separately with age, gender and fluid intelligence used as stratifies.

The main study targeted 123 participants and imaging data were acquired for a random subsample of 78 subjects, 25 of whom were excluded from the analyses in keeping with the image quality criteria (see above). See (43) for details. As a part of the study, half of the subjects received transcranial direct current stimulation (tDCS), whereas the other half received sham stimulation. Since the main study found no effects of stimulation on any of the collected cognitive measures, we did not estimate corresponding effects of stimulation on the imaging data in the present set of analyses, but accounted for stimulation as an additional nuisance regressor during modeling.

### Cognitive training

Between pretest and posttest assessment with cognitive tasks and imaging, a JAVA-based computerized cognitive training battery was administered over 20 sessions (4 weeks, weekends off), each lasting 40 minutes. In the active program, four tasks trained two crucial facets of working memory: updating (running span and n-back tasks) and switching (rule- and task-switching tasks), each with four randomly varying stimuli sets to minimize the use of stimuli-specific strategies. The control intervention was focused on perceptual matching speed, employing four versions of the same task. In the task, participants were asked to assess the similarity of the presented figures and shapes, responding as quickly and accurately as possible. The training batteries were designed to be adaptive, such that the difficulty of the tasks increased systematically with improvements in performance so that the participants always trained at the highest level reached. The batteries were designed and carefully pilot-tested to balance the length and scope of the training, and the working memory training (active) and perceptual matching training (control) groups did not differ in the average level reached by the participants or in overall attendance. The cognitive training took place the week after an extensive 5-day assessment of cognitive functions. See (43) for details and a comparative description of the two batteries.

### Magnetic resonance imaging

Structural and fMRI scans were collected with the GE Discovery MR-750 3.0 Tesla scanner with the 32-channel research head coil located at the Karolinska University Hospital, MRI center in Solna, Sweden. Structural images were acquired employing a standardized contrast-enhanced T1 spoiled gradient (SPGR) “BRAVO” sequence with 0.94 mm^3^ isotropic voxels, FoV 256 mm (256 × 256 matrix), TR/TI/TE = 5.688/450/2.492 ms, flip angle = 12°. Functional MRI data were collected using echo-planar T2* imaging to measure the blood-oxygen-level dependent (BOLD) signal (voxel size 3 mm^3^, TE/TR=30/2000 ms, flip angle = 80°).

### Functional magnetic resonance imaging tasks

Two tasks were administered inside the MR scanner to evaluate connectivity patterns during working memory and reasoning. A figural n-back task assessed working memory performance. Three 1-minute blocks were administered: 1-, 2- and 3-back (each repeated 3 times) with a 4-second fixation screen indicating the rule for the forthcoming level (9:54 min, in total). See **Figure 1** for an illustration. For this task, accuracies were calculated as the average value of true positives and true negatives with a chance level of 50%. The second task was developed to evaluate the functional state involved in complex visuospatial reasoning. Subjects were instructed to complete the presented pattern by selecting one of the four solutions shown at the bottom of the screen. One of the four answers was always correct. Like the n-back task, this paradigm had three levels of difficulty: A, perceptual matching; B, simple analytical (single rule); and C, difficult analytical (with at least two non-trivial rules), similar to the implementation proposed by Yamada et al. (44). Each block was repeated 3 times in a pseudo-random order during the scanning session (9 task-blocks + 9 fixation screens, in total). The reasoning task was self-paced to better match typical fluid intelligence paradigms (such as Raven’s progressive matrices) with fixed (1-minute) blocks (9:54 min of total acquisition time including fixation blocks). The subjects were explicitly instructed not to guess. For the reasoning paradigm, accuracy was defined as fraction of correctly solved problems relative to the total number of attempts per block. In a number of simulations with randomly varying answers and total number of responses, chance-level accuracy was estimated at 25%.

In both tasks, the subjects demonstrated consistent accuracy drop as the difficulty level increased. In all the analyses, performance was measured as total task accuracy, which was defined as the sum of block-specific accuracies.

Both tasks were administered twice: before and after the cognitive training and were programmed and presented using PsychoPy software, version 1.84 (45).

Two tasks were administered inside the MR scanner to evaluate large-scale patterns of brain connectivity underlying working memory and reasoning. A figural n-back task assessed working memory performance. Three 1-minute blocks were administered: 1-, 2- and 3-back (each repeated 3 times) with a 4-second fixation screen indicating the rule for the forthcoming level. The second task was developed to evaluate modifications of the functional connectome involved in complex visuospatial reasoning. Subjects were instructed to complete the presented pattern by selecting one of the four solutions shown at the bottom of the screen. One of the four answers was always correct. Similar to the n-back task, this paradigm had three levels of difficulty: A, perceptual matching; B, simple analytical (single rule); and C, difficult analytical (with at least two non-trivial rules) tasks. Each block was repeated 3 times in a pseudo-random order during the scanning session. The reasoning task was self-paced (to better match typical fluid intelligence paradigms such as Raven’s progressive matrices) within a fixed (1-minute) block. Thus, different subjects completed different numbers of items per block, and the subjects were explicitly instructed not to guess. For the reasoning paradigm, accuracy was defined as the fraction of correctly solved problems relative to the total number of attempts per block.

### Cognitive assessment at pretest and posttest

We evaluated the effects of working memory training via an extensive testing battery completed over four sessions, each lasting about 180 minutes. Pre-intervention testing took place 2 weeks before the training started, and post-testing was completed the week after the intervention. The battery was identical on both occasions and included a number of diverse cognitive tests measuring switching (task- and rule-switching), updating (n-back, running span), attention (temporal expectancy, perceptual matching), episodic memory (verbal and spatial recall), and spatial (WASI, Raven’s, BETA matrices) and verbal (Syllogisms, BIS Analogies, ETS Kit inference) reasoning. See (43) for detailed description of the cognitive assessment.

### Magnetic resonance imaging processing

A standardized SPM12-based (Wellcome Trust Center for Neuroimaging, UCL) pipeline was implemented in the MATLAB R2013a environment (46).

Each subject’s T2* imaging data underwent subsequent steps for slice-timing correction, spatial realignment, and registration to standardized Montreal Neurological Institute (MNI) space using a population-specific template produced from the set of T1 images with the Diffeomorphic Anatomical Registration Through Exponentiated Lie Algebra (DARTEL) algorithm (47).

An extended 24-parameter model (48) was employed for head motion correction. The images were also smoothed (FWHM = 6 mm), de-trended, and, for the connectivity part, also band-pass filtered (0.008–0.12 Hz). The resulting output was used to calculate connectivity matrices (one matrix per each task and load level) based on temporal (Pearson r) correlation between the regions parcellated in accordance with the Craddock’s 200-ROI scheme (49). We did not employ global signal regression due to high risks of removing task-relevant signal variance (see **Supplement, S3**).

No between-task differences were found in mean framewise displacement (meanFD_n-back_ = 0.09±0.04; meanFD_reasoning_ = 0.11±0.09) and neither of the cognitive measures significantly correlated with FD. Nevertheless, we repeated our analyses with scrubbing, which had a very minor impact on the results and did not change any inferences.

After removing negative weights, the adjacency matrices were then used to estimate whole-brain network modularity as implemented in the “Brain Connectivity Toolbox” (50). The Louvain community detection method generalized for weighted graphs was used in the study as a measure of between-network segregation. The method attempts to solve an optimization problem by maximizing modularity (Q) given observed graph weights:

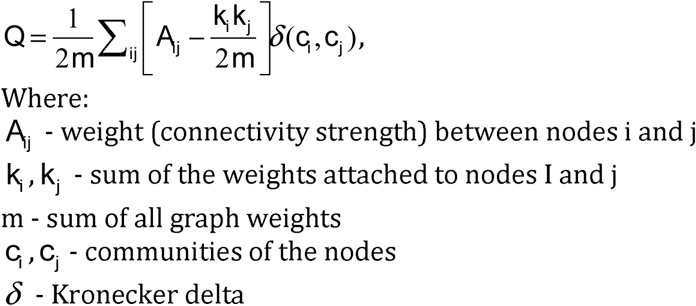

When there are few edges between the communities and within-cluster density is high, Louvain modularity is close to 1, but when the number of within-modular connections is comparable to the one from a random graph, modularity usually approaches 0. Practically, it measures level of modular segregation in a graph.

### Statistical analyses

After the preprocessing and cleaning, behavioral and imaging measures underwent outlier detection and checks for distribution normality. Between-group comparisons of demographic data used two-sample t-tests for continuous variables and binomial tests for sex ratios.

Next, we conducted a number of analyses to evaluate the reliability and validity of the collected measures. Pre-post Pearson correlation coefficients were calculated for the behavioral data to evaluate reliability. To address the problem of task validity, we analyzed updating and spatial reasoning domains in latent space, employing structural equation modeling (SEM) of the two constructs with *offline* measures and data collected inside the MRI scanner (See **Figure 2**). Each of the *offline* latent variables (LVs) was modeled with 3 tasks (updating: n-back and two running span tasks; spatial reasoning: Raven’s progressive matrices, WASI, and BETA). *Online* (fMRI) LVs were formed with two indicators each (updating: 2-back and 3-back; spatial reasoning: “simple analytical” and “difficult analytical”; 1-back and reasoning “A” accuracies were excluded from the SEM due to ceiling effects). The variance of the LVs was constrained to 1 to make path coefficients equivalent to Pearson correlations. SEM was also repeated for block-specific measures of in-scanner performance (2-back and “simple analytical”; 3-back and “difficult analytical”).

**Fig. 2.**
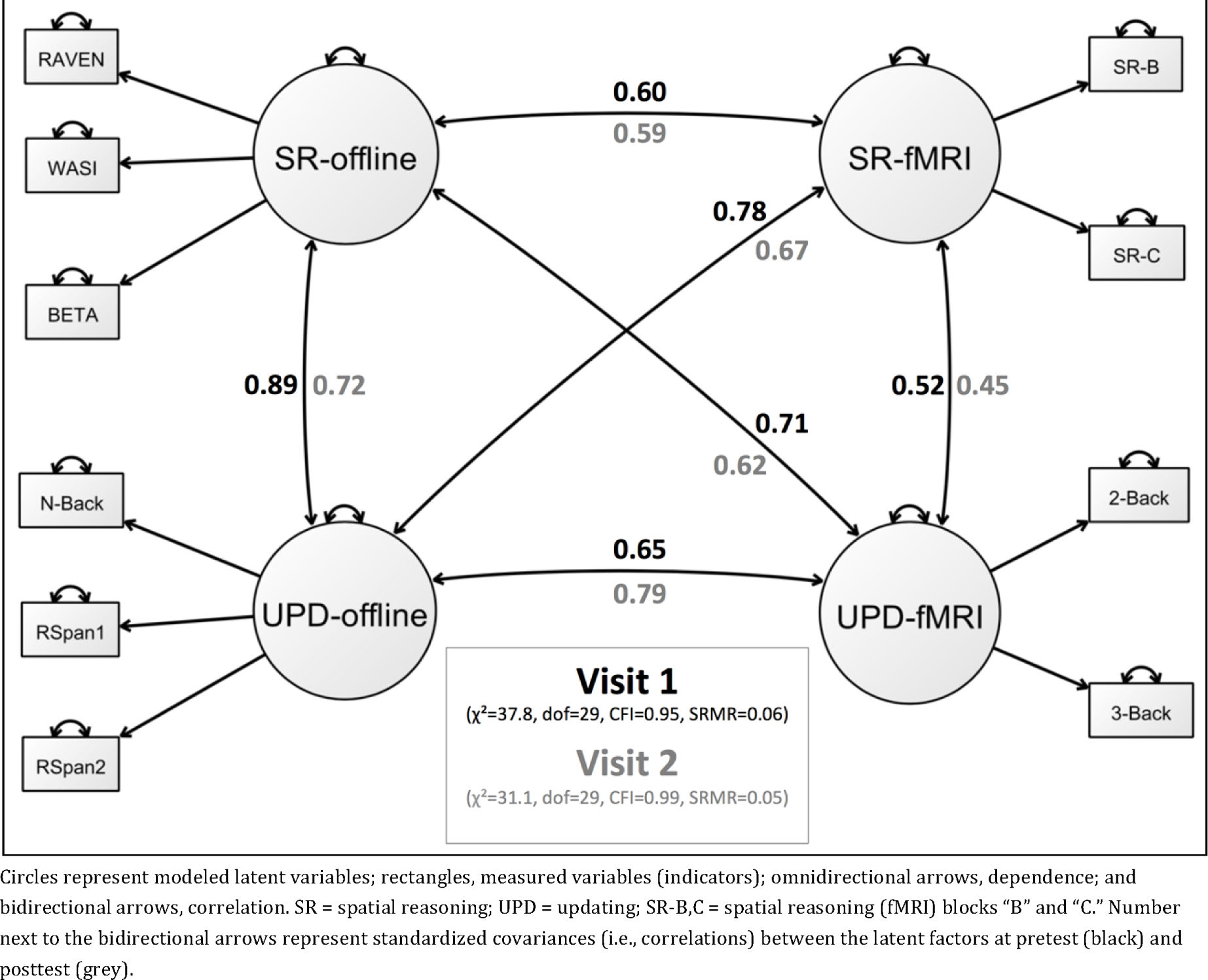
Results of the structural equation modeling analysis.

The next block was focused on evaluating task-related BOLD activity and modularity at baseline. First, we estimated first-level contrasts in both tasks: a) 2-back – 1-back, b) 3-back – 1-back, c) reasoning “B” – reasoning “A,” and d) reasoning “C” – reasoning “A.” Then we performed a second-level analysis of the activations, which yielded block-specific statistical maps. Once the smoothness of the data had been estimated, adjustment for multiple tests was carried out using Monte Carlo simulations (n=5000) as implemented in the AlphaSim algorithm (51) with an initial cluster-forming threshold of p<0.005. In addition, individual block-specific activation patterns were matched with a meta-analytical map derived using Neurosynth software (http://www.neurosynth.org/) (52). This procedure employs voxel-wise Pearson correlation as a measure of similarity (53) between the statistical map and the meta-analytical image generated for a specific feature of interest. In the present analysis, the search was completed for the terms “working memory” (n=901 studies) and “reasoning” (n=147). The search was also completed for “executive control” (n=157), which was used as a third independent term to evaluate validity of the activation patterns.

Next, we embarked on the analyses of large-scale connectivity, evaluating the differences in functional connectome between the tasks and between-task correlation of modularity across different loads. Subsequently, we conducted a number of mass-univariate tests of raw functional connectivity to identify which of the networks and between-network interactions were involved in the reported global effects. This was accomplished by fitting a linear model that evaluated correlations between block-specific functional connectivity and total task performance.

The last analytical block addressed effects of working memory training on performance and large-scale functional connectome. We started by evaluating the main effects of time on task performance and then estimated load × group × time effects in each of the two tasks using linear mixed-effects models (random effect: “subjects”). Standardized effect sizes were also calculated and reported as the mean difference in performance divided by the pre-test standard deviation.

Modularity was analyzed as overall pre/post change, as well as Time × Group interactions. At this stage, a full-interaction model was fitted to evaluate the effects of working memory training in two tasks (modularity ~ load × task × time × group × WM performance, random effect: “subjects”). Finally, a subsequent analysis of raw functional connectivity was conducted to localize the reported global effects by estimating Group × Time effects on connectivity for each block.

### Statistical software and toolboxes

All statistical analyses of behavioral data and modularity were performed with the R programming language, version 3.2.2 (54) and “nlme” toolbox (55). In all models, age, sex, and first 3 principal components from the motion parameters were introduced as nuisance covariates.

Mass-univariate analyses of BOLD activation patterns were carried-out in SPM12 (Wellcome Trust Center for Neuroimaging, UCL).

The Network-Based Statistics Toolbox (56) was used to analyze raw functional connectivity.

Structural equation modeling was conducted using Ωnyx software (57).

### Data availability

Raw MRI and behavioral data can be requested from the authors and transferred for specific analysis projects that are in line with the original ethical approval. This requires a data use agreement, which effectively transfers the confidentiality obligations of the institution (Karolinska Institutet) at which the original research was conducted to the institution of the recipient of the data. An R-script specifying main analytical and plotting steps is available at: https://github.com/alex-lebedev/RBT1_modularity

## Results

### Cognitive measures

Initial analyses aimed to probe the reliability and validity of the cognitive tasks. Between-person differences in performance on the in-scanner tasks **(Figure 1**) were relatively stable from before to after cognitive training, with pre-post correlations of *r*= 0.57 (p<0.001) for n-back and *r*=0.6 (p<0.001) for total reasoning performances averaged across the difficulty levels of the tasks. Corresponding values were equivalent between the groups: r_pre-post_=0.54/0.47 (z=0.32, p=0.75) in control/active groups for n-back and r_pre-post_=0.57/0.65 (z=-0.44, p=0.66) in control/active groups for the reasoning task. These results indicate satisfactory lower bounds of reliability.

The validity of the in-scanner tasks was evaluated with structural equation modeling (see **Methods**). The n-back task performed in the scanner (UPD-fMRI) correlated strongly (pre: *r =* 0.65 p<0.001; post: *r* = 0.79, p<0.001) with a latent factor of working-memory updating (formed by tasks assessed outside the scanner; see **Figure 2** and **Methods** for a more detailed description of the testing battery). Performance on reasoning task administered in the scanner also correlated strongly (pre: *r* = 0.6, p<0.001; post: *r* = 59, p<0.001) with a reasoning factor (formed by three tasks assessed outside the scanner). Cross-correlations were equivalent. Performance on the in-scanner n-back correlated in a statistically significant way with the reasoning factor (pre: *r* = 0.71, p <0.001; post: *r* = 0.67, p <0.001), and performance on the in-scanner reasoning task, with the working-memory updating factor (pre: *r* = 0.78, p<0.001; post: *r* = 0.67, p<0.001). The reasoning and working-memory updating factors were highly correlated (pre: *r* = 0.89, p<0.001; post: *r* = 0.72, p<0.001), and so were the in-scanner tasks (pre: *r* = 0.52, p<0.001; post: *r* = 0.45, p<0.001). Overall, the reliability as well as discriminant and convergent validity of the tasks used in the scanner were thus satisfactory.

### Task-related BOLD activity and modularity at baseline

Both tasks produced similar and reliable load-related activations in the control network (see **Figure 3**) that are typical for working memory and reasoning tasks. Moreover, similarity (voxel-wise Pearson r, see **Methods** for details) of individual activation patterns to the meta-analytical map derived using the Neurosynth package (see **Figure S1**) for “executive control” was statistically significant across all loads (significantly larger than 0 at p<0.001) and positively correlated with subjects’ performance during n-back (2-back: r=0.34,p=0.01; 3-back: r=0.43, p=0.001) and simple analytical reasoning tasks (reasoning-B: r=0.30, p=0.03; reasoning-C: r=0.19, p=0.18). These results indicate that both paradigms reliably engaged the executive control circuit and that the magnitude of the engagement significantly correlated with successful task solving in all active conditions except for the difficult analytical block (“C”) of the reasoning task, in which engagement of the circuit had a limited non-significant effect on performance.

**Fig. 3.**
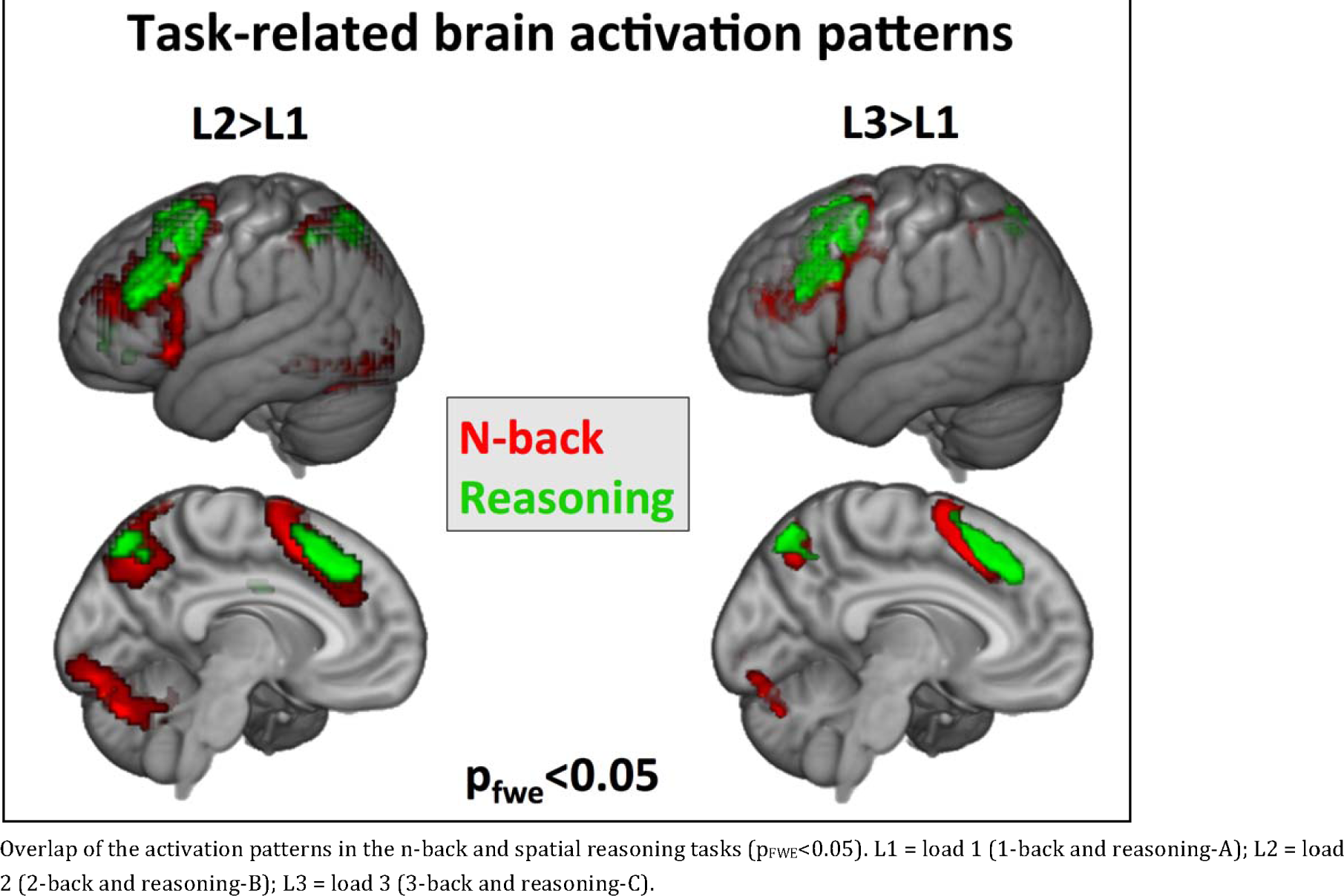
Overlap of the task-related activation patterns

Despite similarities in the activation patterns, we observed a statistically significant effect of load, t(208)=-2.7, p<0.01 and load-by-task effect, t(208)=2.74, p=0.007 on modularity (Q). These effects were further qualified by a load-by-task-by-performance effect, t(208)=3.07, p=0.002. Although performance was implemented as a continuous variable in these analyses, to visualize these effects in an accessible manner we performed a median split on performance **(Figure 4**). The resulting graph shows that individuals who performed well on the n-back task had a tendency to increase modularity (i.e., between-network segregation) with load, whereas individuals with better performance in the more complex reasoning task down-regulated modularity with task complexity. In other words, superior performance in the difficult working memory task (n-back) was associated with higher modularity (between-network segregation), whereas superior performance in the complex reasoning task was associated with lower modularity (see also **Figure 5A**). To further assess these effects, we selected a subset of subjects (n=17) who performed high on both tests, defined as above-median performance both in n-back and reasoning. Load-by-task effect was significant in this group, t(82)=3.28, p=0.001, whereas subjects performing poorly on both tasks (n=18) did not exhibit such pattern, t(87)=3.28, p=0.001. Thus, individual differences in performance were, as predicted, related to different (segregated vs. integrated) configurations of the functional connectome in the working memory and complex reasoning tasks, especially at higher loads.

**Fig. 4.**
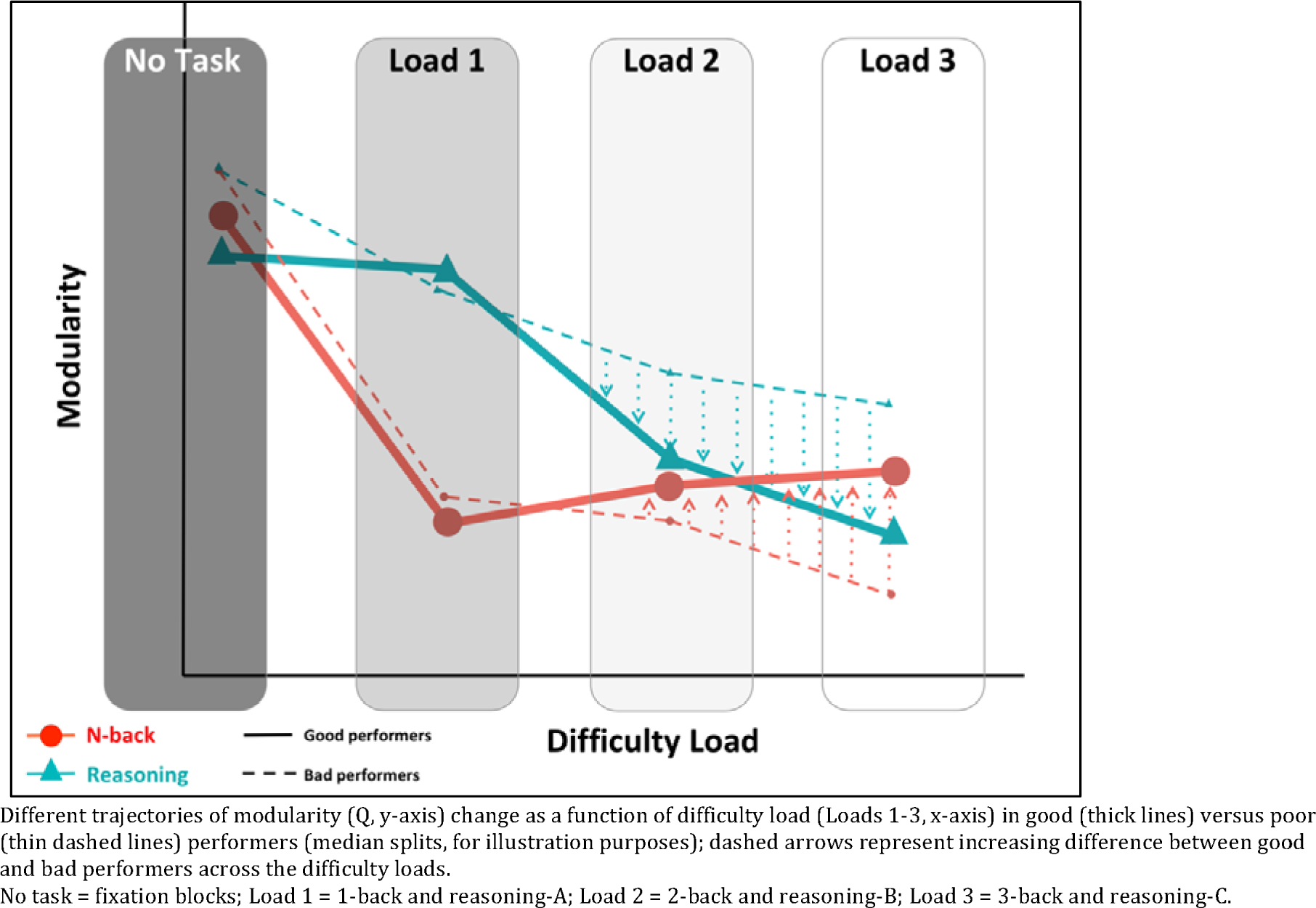
Shifts in modularity associated with successful solving of working memory and reasoning tasks.

**Fig. 5.**
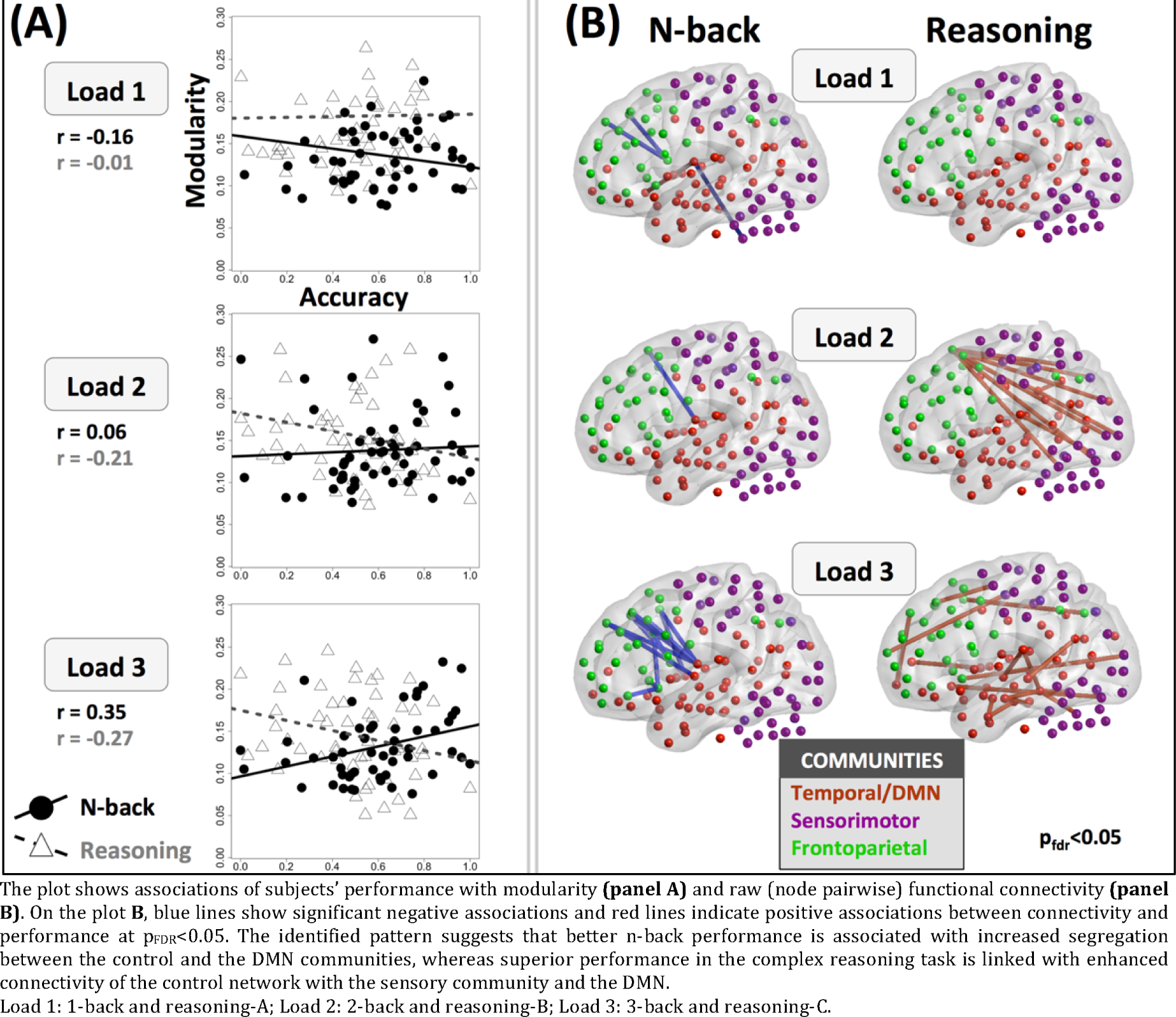
Different associations between large-scale functional connectome and performance in n-back and reasoning tasks.

To identify which of the networks and between-network interactions drive these relations with global modularity, we conducted a number of subsequent tests of raw functional connectivity (56). The results presented a similar pattern of findings, which suggest that superior n-back performance is linked to increased segregation between the control and the DMN networks, whereas superior performance in the complex reasoning task is linked with more active cross-talk of the control network with the sensory community and the DMN **(Figure 5B**).

Note that modularity during the fixation condition did not differ in a statistically significant way between the tasks, t(52)=0.34, p=0.736. Moreover, both tasks induced prominently less modular states than the fixation condition: N-back, t(52)=6.2, p<0.001; reasoning, t(52)=7.0, p<0.001 **(Figure 4**). Yet, all difficulty loads were consistently associated with modular states: mean modularity was significantly (p<0.001) and consistently larger than the one derived from a set (n=250) of simulated random networks with equivalent properties (see **Supplement, S2**), both as a total effect and when taken separately within specific tasks and loads.

### Effects of working memory training on performance and modularity

The main effect of time (pretest vs. posttest) on performance was statistically significant, which indicated general improvements in n-back and reasoning performance (0.57 SDs for n-back, t(52)=4.97, p<0.001; 0.64 SDs for the reasoning task, t(52)=4.02, p<0.001; **Figure 6**). Working memory training led to larger increases in n-back performance at higher loads than perceptual matching training did (load-by-group-by-visit: t(155)=2.77, p<0.01). T-tests comparing gains from working memory training with gains from perceptual matching training at different loads indicated a statistically significant effect at 3-back only. There was a 0.88 SD improvement in the working memory training group and a 0.1 SD improvement in the control group, t(43.029)=2.92, p=0.003. Neither the group-by-visit nor the load-by-group-by-visit effects on reasoning performance were statistically significant, which confirmed the lack of far transfer effects of working memory training to reasoning published in our previous report of the behavioral data from this study (43). Subsequent t-tests revealed that improvement in block “B” accuracy (simple analytical) tended to be larger in the active (0.695 SD) than the control group (0.13 SD), t(46.331)=-2.01, p=0.026. It is worth noting, however, that this comparison would not survive strict p-threshold correction for multiple testing (0.05/3 tests = 0.016).

**Fig. 6.**
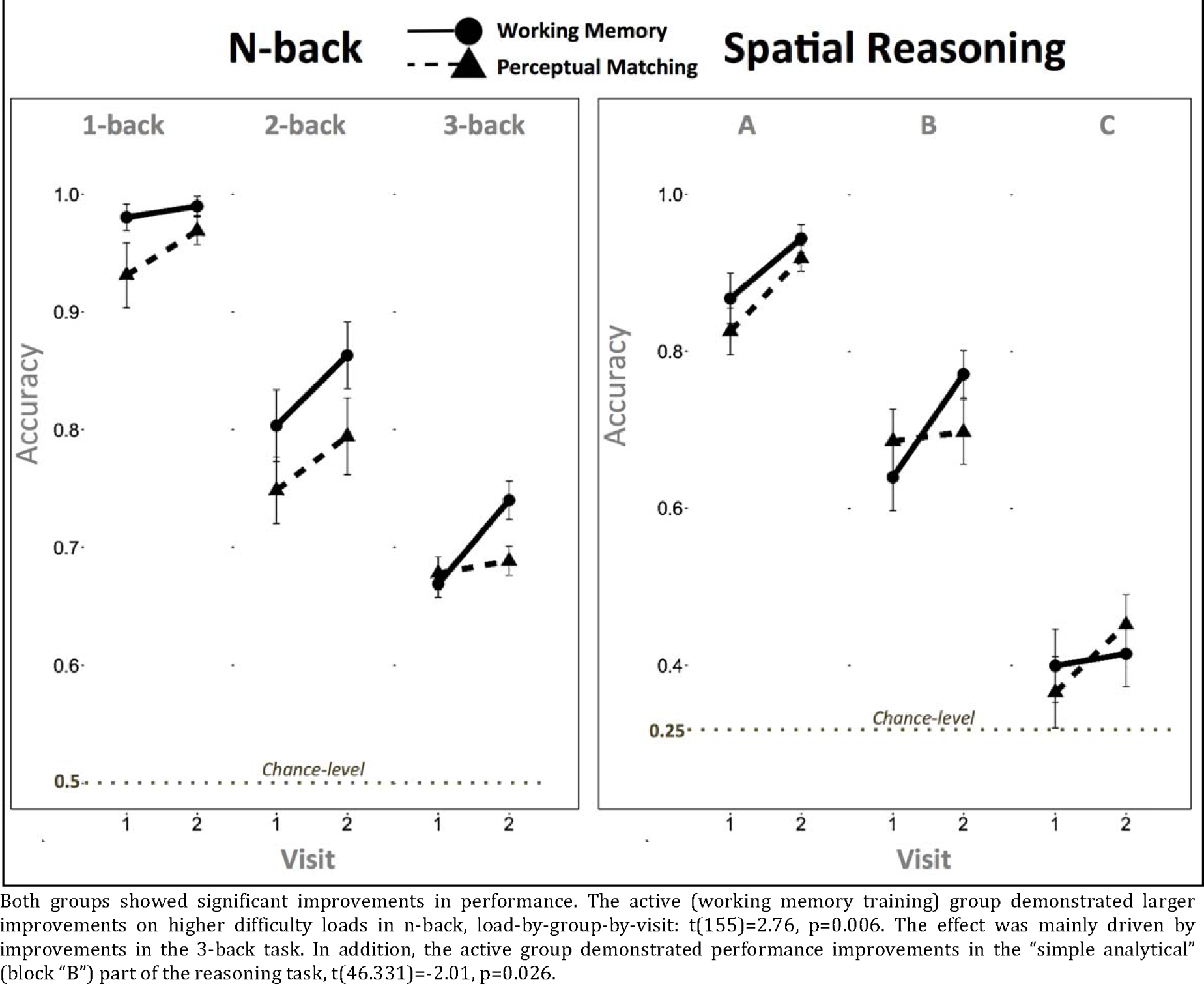
Effects of working memory training on in-scanner n-back and reasoning

We found that the active and control interventions affected modularity differently, with a statistically significant group by time interaction, t(316)=3.5, p<0.001 **(Figure 7**). Taken separately, working memory training resulted in a significant increase in modularity, 0.31 SD, t(161)=3.3, p=0.001, whereas perceptual matching training did not result in statistically significant changes in modularity. Instead, it exhibited a trend toward reducing it, -0.2 SD, t(155)=-1.8, p=0.07. The group-by-load-by-time effect, however, was non-significant, which is consistent with a load-invariant effect of working memory training on modularity. The group-by-task-by-time interaction was statistically significant t(310)=2.25, p=0.025, indicating that effects of training on modularity were somewhat larger for the reasoning task.

**Fig. 7.**
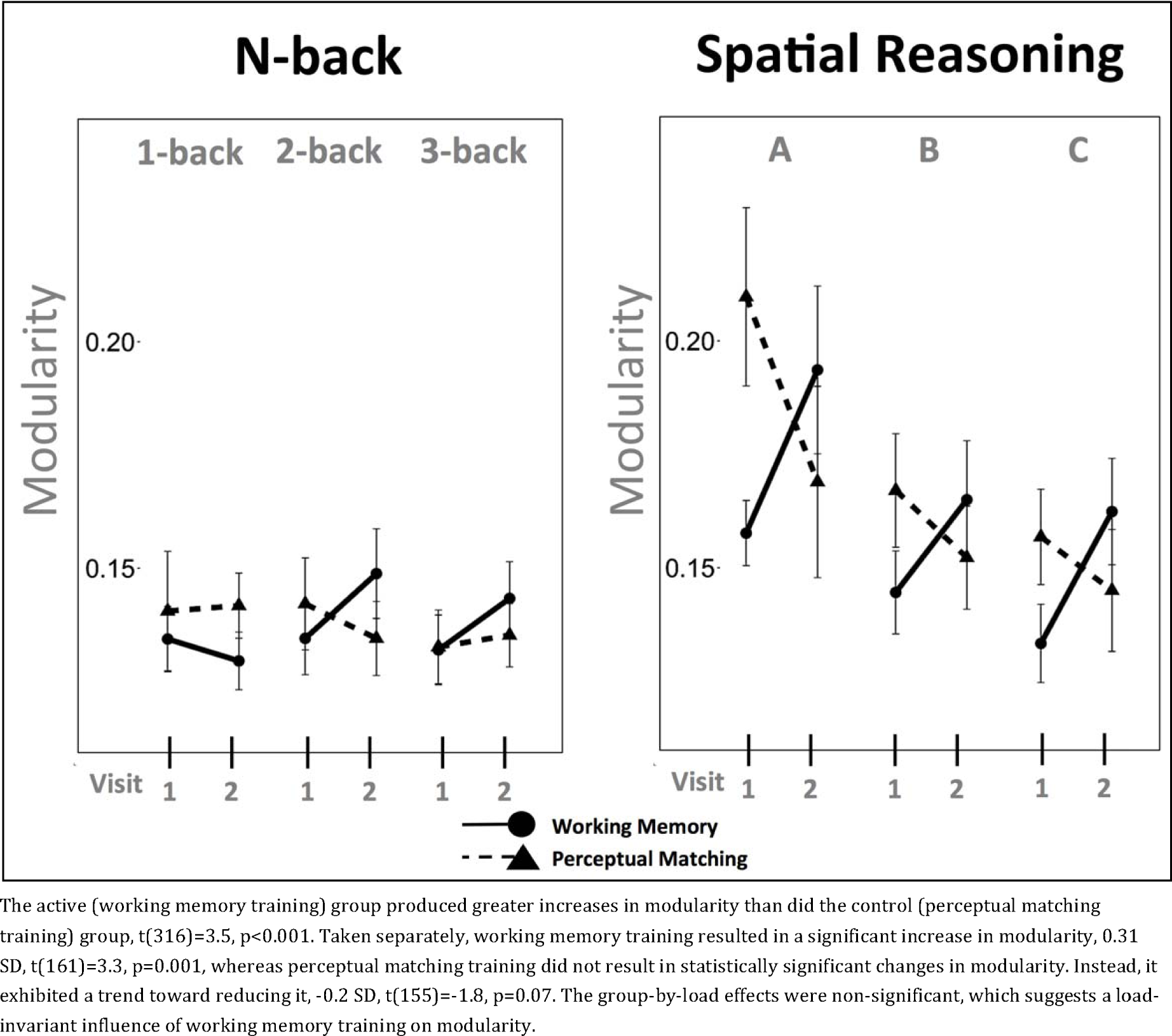
Effects of working memory training on modularity

Longitudinal analyses of raw functional connectivity revealed that working memory training reduced connectivity of the frontoparietal network mostly with the sensorimotor community and some components of the default mode / temporal cluster, with more marked effects seen in the reasoning paradigm (Group × Time T-contrast: p_fdr_<0.05, **Figure 8**).

**Fig. 8.**
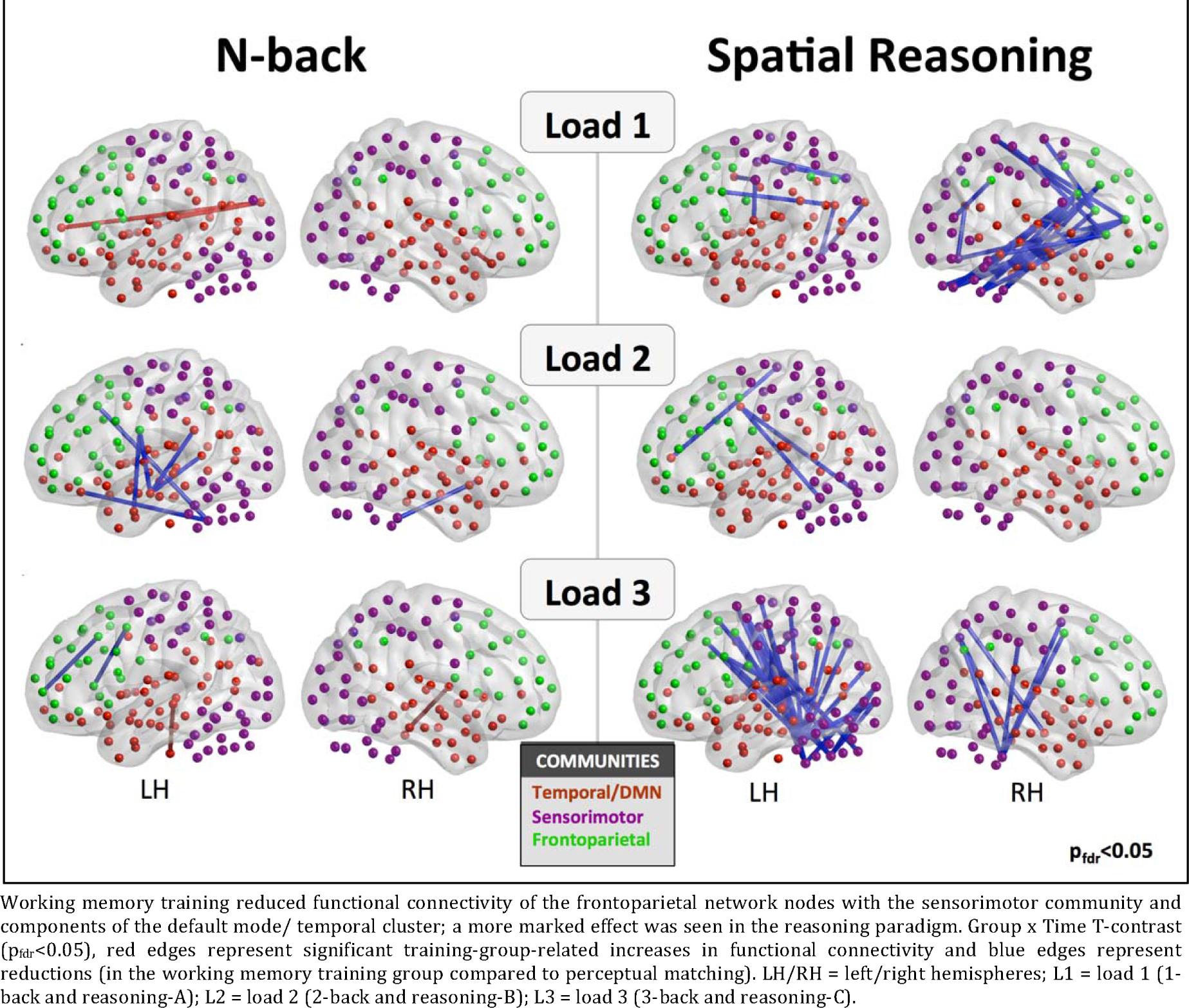
Effects of working memory training on functional connectivity

Given that the two tasks benefited from opposite modes of large-scale brain dynamics whilst training intervention resulted in load-invariant increases in modularity, we performed a set of additional analyses to explore whether individual differences in training-induced changes in working memory performance and large-scale functional organization were related to changes in to reasoning performance. To do this, we first fit a simple model using behavioral data collected offline and observed that overall gains were negatively related between working memory (trained) and reasoning (transfer) compoaites [r=-0.28, ΔSR_composite_~ ΔWM_composite_: t(51)=-2.01, p=0.04], and this relationship was trend-significant after the main effect of TrainingGroup had been removed [r=-0.26, t(51)=-1.95, p=0.06]. For the in-scanner performances, however, a corresponding effect was non-significant [r=0.17, t(51)=1.3, p = 0.2].

Subsequently, exploring whether training-induced increases in modularity during reasoning task had negative relationships with changes in reasoning performance did not yield any significant results for any of the difficulty loads. However, we found that change-change correlations between modularity during n-back task and reasoning performance gradually went down across the three loads (r_Load1_=0.19, r_Load2_=0.07, r_Load3_=-0.15), but this was a trend that did not reach conventional levels of statistical significance, as shown be the Load-by-Modularity effect on performance (t(158)=1.75,p=0.08).

## Discussion

Employing methods of large-scale network analyses, we found that despite high correlations in the behavioral data and substantial overlap of brain activation patterns, performance in complex reasoning and working memory tasks was associated with different modes of large-scale connectivity. Better performance in a typical working memory task (n-back) was associated with higher modularity (i.e., increased between-network segregation), whereas better performance in a reasoning task was associated with lower modularity (i.e., more interactions between the major brain networks) at higher task-complexity. Furthermore, working memory training modified functional connectome in both tasks in a similar way; that is, by increasing modularity. Perceptual matching training, on the other hand, did not have any significant impact on it. The observed effects appeared to be driven by integration within the frontoparietal control network and its segregation from the other communities.

It has been proposed that balanced modular organization of the brain is necessary for the emergence of the critical dynamics important for adaptive information processing and learning (18-20, 58-60). Previous research has indicated that most cognitive tasks generally tend to perturb these modular dynamics by shifting them to a more integrated state of increased between-network communication (24, 29-31, 61, 62). We observed a similar pattern of general modularity reductions from rest to task in the present study. However, we also showed that performance on working memory and complex reasoning tasks is associated with modularity in different ways, such that between-network interplay benefits complex reasoning, whereas sustaining a more modular organization benefits working memory performance.

The observed divergent pattern of shifts in connectivity is generally in line with the literature, which suggests that automated tasks requiring outward attentional focus tend to favor segregation of the DMN and the frontoparietal CN (34, 35, 63), whereas tasks requiring novel and creative problem solving benefit from enhanced between-network interplay (13, 36). It may be that lower modularity is associated with better complex reasoning performance because it relates to an exploratory mode of cognition (39) beneficial for rule discovery and abstraction (41). In contrast, such a mode of cognition may hinder working memory performance, which relies on a more modular network configuration with stabilized dynamics of the frontoparietal network coupled with effective suppression of DMN activity. These distinct associations were more marked at higher task complexity. The findings fully support the network neuroscience theory of human intelligence suggesting that the tasks that engage broader cognitive functions benefit from dynamic shifts in connectivity patterns toward a more random reconfiguration, unlike the ones relying on specific cognitive abilities that rather benefit from a more regular mode of large-scale organization of the connectome (60). This possibility is also in line with the recent findings that performance on simple tasks is positively related to resting state modularity, whereas performance on complex tasks is negatively related to resting state modularity (38).

It is worth noting that the circuit described in the influential parieto-frontal integration theory (P-FIT) of intelligence represents a broader community than the frontoparietal network discussed in the present study and also includes components of the default mode and visual clusters (64). The reported results thus do not contradict P-FIT. Instead, our observation that superior reasoning performance favors enhanced cross-talk of the frontoparietal regions with occipital and medial prefrontal cortices is nicely in line with the P-FIT model.

Working memory training produced marked increases in global modularity, thus promoting the pattern of large-scale dynamics that was associated with better working memory performance prior to training. Interestingly, the same pattern of modularity changes was observed for the reasoning task although no improvements in total reasoning performance were observed. This suggests that working memory training fosters a mode of solving the reasoning tasks that relies more on working memory and less on an exploratory mode of cognition. Such an interpretation is consistent with the trends for training-related improvements on the “simple analytical” (B) reasoning tasks, which may benefit from superior modularity (because the task demands attention, but the rules for solving it are simple to detect and to abstract) but not on the “complex analytical” (C) tasks, which may benefit from less modular dynamics (because they require complex pattern discovery). It should be acknowledged that the ability to solve the complex analytical blocks is generally more representative of fluid intelligence. Therefore, despite potential trends in training-related differences in performance improvements on the simple analytical part of our reasoning task, our study does not support the thesis that it is possible to improve the spatial aspect of fluid intelligence with working memory training, at least in healthy older people. This conclusion is generally in line with the literature, which shows that working memory training has either no effects at all (65) or, at best, small effects (66) on reasoning. Our results expand these observations: they suggest that the reason working-memory training does not improve fluid intelligence is because it shifts large-scale brain dynamics in ways that are not beneficial for solving novel and complex reasoning problems.

In summary, although inter-individual differences in reasoning and working memory performance, and most other cognitive tasks too, are positively correlated, the brain may solve these tasks in partly different ways. Rather than indicating strong links between the processes involved in reasoning and working memory, the correlated inter-individual difference could be produced by initially unrelated cognitive domains that support each other in learning and development over the lifespan (67). Indeed, it is clear that higher working memory capacity does not necessarily imply superior intelligence. A particularly illustrative example is the observation that chimpanzees have, in fact, better visual working memory than humans (68), which suggests that the cognitive advancement of the human species was unlikely driven by increased storage capacity. Despite the fact that our study did not find that changes in modularity induced by working memory training to actually be detrimental to reasoning abilities, it generally supports the idea that storage capacity may, in fact, compete with some aspects of complex reasoning, such as insight and deep information processing capabilities (69). Our results are also broadly in line with the recently proposed idea of a trade-off in biological systems, in which high-modular organization provides evolutionary benefits over shorter timescales, and low-modular systems exhibit greater fitness over longer timescales (38, 70). Notably, imposing time constraints on reasoning problems appear to increase the correlation between subjects’ performance on such tasks with their performance on working memory tests (4). The trade-off hypothesis (38, 70) and our results are in accordance with the observation that working memory training studies tend to show transfer effects when reasoning tasks are administered under more stringent time constrains (71) (possibly benefiting from high-modular states), whereas those that administer the tasks in a traditional, time-relaxed, manner (that benefit from low-modular dynamics) do not report such effects (72).

This study had a number of limitations. First, one could argue that the tasks we used inside the scanner do not properly represent corresponding abilities of working memory and fluid intelligence, or at least that the tasks only tap into selected aspects of these abilities. For example, some tasks that are supposed to characterize the same domain of working memory (e.g., complex span and n-back) actually have relatively low correlation and may therefore represent different aspects of working memory (73). Meanwhile, some researchers have questioned the plausibility of this argument, as several studies demonstrate that the performances on the two tasks converge when working memory is modeled as a latent construct (74, 75). This is still of crucial importance for the present study, considering that both resting state and task-related modularity has been shown to be differently associated performance on working memory tasks, depending on their complexity with positive correlation found for visual working memory performance and a trend-level negative correlation observed for a complex span task (76).

Another potential limitation of the study is the absence of counter-balancing of the fMRI tasks, which may have affected the reliability of the results because of carry-over effects of the n-back paradigm on reasoning. However, consistent engagement of the control circuit in both tasks, strong associations between modularity and performance, together with the occurrence of longitudinal changes in the expected direction strongly support plausibility of the proposed interpretations. It is also worth noting that even though we tried to minimize the difference between the tasks in aspects that are not central to the research question, the stimuli still differed in a number of ways, besides in their reasoning and working memory demands, which, in turn, might confound some of our results. Finally, the present study was conducted on a sample of older adults, and the results therefore may not be generalizable to younger populations, especially since community structure of the brain undergoes substantial reconfiguration over different periods of the lifespan (21, 22). Decreased signal-to-noise ratio of brain activity found in older subjects (77) may foster a different mechanism underlying rest-to-task perturbation of functional connectome in this population, perhaps imposing higher demands on the aging brain to upregulate segregation of the frontoparietal network to allow superior filtering of the neural noise, which in turn appears to have detrimental effects on working memory capacity (78), whilst facilitating exploratory states of brain dynamics (79). Further studies may want to focus on how differences in the amount of neural noise influence reconfiguration of the functional connectome depending on task complexity. This, in turn, may help to explain a discrepancy of findings from imaging studies of working memory, some of which show positive associations between modularity and performance (23), whereas others report negative relationships (62).

Despite these issues, we conclude that reasoning performance and working memory performance in older adults have different associations with large-scale functional brain connectivity at higher task complexity. We also found that working memory training fosters task-invariant increases in functional network modularity and does not improve performance on complex reasoning tasks. Instead, complex reasoning tasks benefit from a higher degree of between-network integration. Together, these results indicate that complex human reasoning abilities cannot be reduced to working memory and add weight to the conclusion that working memory training interventions will not result in improvements that transfer to broader cognition.

## Acknowledgments

We thank Anders Rydström, Linda Lidborg, Jakob Norgren, Helena Franzén, and Simon Peyda for help with data collection and Marie Helsing for help with recruitment and organization. Thanks also to Alireza Salami, Goran Papenberg and Lars Nyberg for comments on an earlier draft of this article. This research has received funding from the European Research Council under the European Union’s Seventh Framework Programme (FP7/2007-2013) / ERC Grant agreement # 617280 - REBOOT. Martin Lövdén was also supported by a “distinguished younger researcher” grant from the Swedish Research Council (446-2013-7189).

